# PRMT5 deficiency disturbs Nur77 methylation to inhibit endometrial stromal cell differentiation in recurrent implantation failure

**DOI:** 10.1101/2024.02.06.579055

**Authors:** Zhiwen Cao, Xiaoying Wang, Yang Liu, Xinyi Tang, Min Wu, Xin Zhen, Nannan Kang, Lijun Ding, Jianxin Sun, Xinyu Cai, Haixiang Sun, Guijun Yan, Ruiwei Jiang

## Abstract

Various posttranslational modifications (PTMs) have been implicated in endometrial stromal cell (EnSC) differentiation, but the potential role of PTM crosstalk has not been identified. Here, we report that protein arginine methyltransferase 5 (PRMT5) is indispensable for human endometrial decidualization, functioning as a key regulator of decidualization defect in recurrent implantation failure (RIF) patients. Uterine-selective deletion of *Prmt5* led to defective embryo implantation in mice due to impaired EnSC decidualization. Mechanistically, we find that PRMT5 catalyzes symmetric dimethylation of orphan nuclear receptor Nur77 at arginine 346, which in turn promotes Nur77 nuclear localization and increases its transcriptional activity in EnSC. Moreover, we demonstrate that PRMT5-mediated Nur77 methylation antagonizes AKT-induced phosphorylation of Nur77 at serine 351 in the transition from proliferation to differentiation of EnSC and disruption of the balance between methylation and phosphorylation of Nur77 is essentially involved in the endometrium of RIF patients. Furthermore, by modulating the methylation-phosphorylation of Nur77 and its transcriptional activity, we rescued impaired decidualization in RIF, further highlighting the critical role of the PRMT5/AKT/Nur77 complex in uterine receptivity to embryo implantation.

## Introduction

Reproduction health has been emerging as an important and challenging issue around the world. A variety of social, educational, environmental and lifestyle pressures associated with modern affluent society contribute to a severe decline in the total fertility rate(1). Meanwhile, widespread infertility and the need for assisted reproductive technologies (ART) are now major health issues(2). It’s reported that about 12.7% of reproductive age women seek infertility treatment each year in the US, and more than 1 million assisted reproduction technology treatment cycles in 2018 have been reported for the first time in Europe(3, 4). Success rates of ART have improved dramatically since the first live birth from in vitro fertilization in 1978. However, good quality embryos transferred to an anatomically normal uterus still fail to implant in some women, even after three attempts, which was defined as recurrent implantation failure (RIF)(5, 6). The ability of the endometrium to allow normal implantation is termed receptivity, while two-thirds of implantation failures are secondary to suboptimal endometrial receptivity(7).

Fertile women attain normal endometrial receptivity during the mid-luteal phase (days 21–24 of a 28-day cycle), named as “window of implantation”, which was driven by estrogen, progesterone and downstream molecular responses(8). Decidualization is characterized by the decidua transformation of endometrial stromal cells, differentiating from elongated fibroblast-like cells into rounded or polygonal-shaped decidual cells, in which large numbers of secretory proteins, including prolactin (PRL) and insulin-like growth factor binding protein-1 (IGFBP1), two well-known decidualization markers, were produced. It is important to understand the key decidual pathways that orchestrate the proper development of the placenta. Besides the key role of progesterone and progesterone receptor (PR), many important transcriptional factors and signaling transduction pathways, including Forkhead Box O1 (FOXO1), CCAAT/enhancer-binding protein β (CEBPB), HOXA10, Nur77, cyclic AMP (cAMP) signaling, signal transducers and activators of transcription (STAT) signaling, AKT signaling, and TGFβ signaling, have been reported to contribute to the establishment of decidualization(9). Decidualization constructs a soft spongy substance as a suitable support matrix for successful implantation, whereas decidual defects have been revealed by our and other groups to be associated with RIF in recent years(10–14).

Posttranslational modifications (PTMs) of proteins are essentially implicated in the dynamic and reversible alterations of various protein properties and functions. As the key upstream regulator in decidualization, PR has been reported to be modified by phosphorylation, ubiquitination, acetylation, SUMOylation and methylation(15–17). As one of the core transcription factors for decidualization, FOXO1 was subjected to phosphorylation, ubiquitination and acetylation(18). The crosstalk between these different modifications further increases the proteome complexity(19). While PTM crosstalk has been studied at depth in many biological processes, its roles in decidualization remain less explored. Recently, protein arginine methylation, as catalyzed by protein arginine methyltransferases (PRMTs), has emerged as an important PTM implicated in the regulation of many biological processes, including DNA repair, RNA processing, chromatin regulation and signal transduction(20). Interestingly, protein arginine methylation has been shown to vigorously interact with other PTMs, such as phosphorylation and ubiquitination (21–23). Our previous study has preliminarily found that PRMT5 is involved in the decidualization *in vitro*, but the detailed mechanism is not yet known (24).

Here, we reported that stromal PRMT5 is indispensable for endometrial decidualization in mice and humans through interacting with orphan nuclear receptor Nur77. Additionally, we found that PRMT5-mediated methylation of Nur77 at R346 augmented Nur77 transcriptional activity through attenuating AKT-mediated Nur77 phosphorylation at S351. Furthermore, tipping the balance of these two modifications toward methylation can efficiently improve the decidualization defect. In this regard, our results provide novel regulatory mechanism of PRMT5/AKT/Nur77 underlying regulation of endometrial stromal cells proliferation and differentiation during the embryo implantation.

## Results

### Loss of PRMT5 in endometrial stromal cells leads to embryo implantation failure and decidualization defect in mice

To gain insight into the expression pattern of PRMT5 in early pregnancy, we performed immunohistochemistry in the uterus of mice from days 1-8 of pregnancy. As shown in Supplemental Figure 1A, PRMT5 expression was weak in the pre-implantation uterus on day 1 and mildly increased in the peri-implantation uterus on day 4. After embryo implantation, PRMT5 was moderately upregulated in the stromal cells surrounding the blastocyst on day 5, while it appeared more abundant in the decidualizing cells on day 6 and day 8. We generated the mouse model with conditional ablation of PRMT5 in the endometrial stroma using anti-Mullerian hormone type 2 receptor *(Amhr2)-Cre* (*Amhr2^cre^Prmt5^f/f^*). The knockout efficiency was determined by western blot and qRT-PCR assays of the uteri from the control and *Amhr2^cre^Prmt5^f/f^* mice (Figure 1A, and Supplemental Figure 1B). In the *Amhr2^cre^Prmt5^f/f^* uteri, PRMT5 protein was virtually ablated in the stromal cells of the uterine anti-mesometrial pole, whereas the expression of PRMT5 remained intact in the epithelial cell layer as well as in the stromal cells in the mesometrial pole, which was consistent with expression of the Cre-recombinase in the *Amhr2^cre^* mouse line (Figure 1B)(25). Furthermore, the primary endometrial stromal cells (EnSCs) isolated from *Amhr2^cre^Prmt5^f/f^*mice showed reduced expression of PRMT5 than that from *Prmt5^f/f^*mice (Figure 1C).

**Figure 1.**
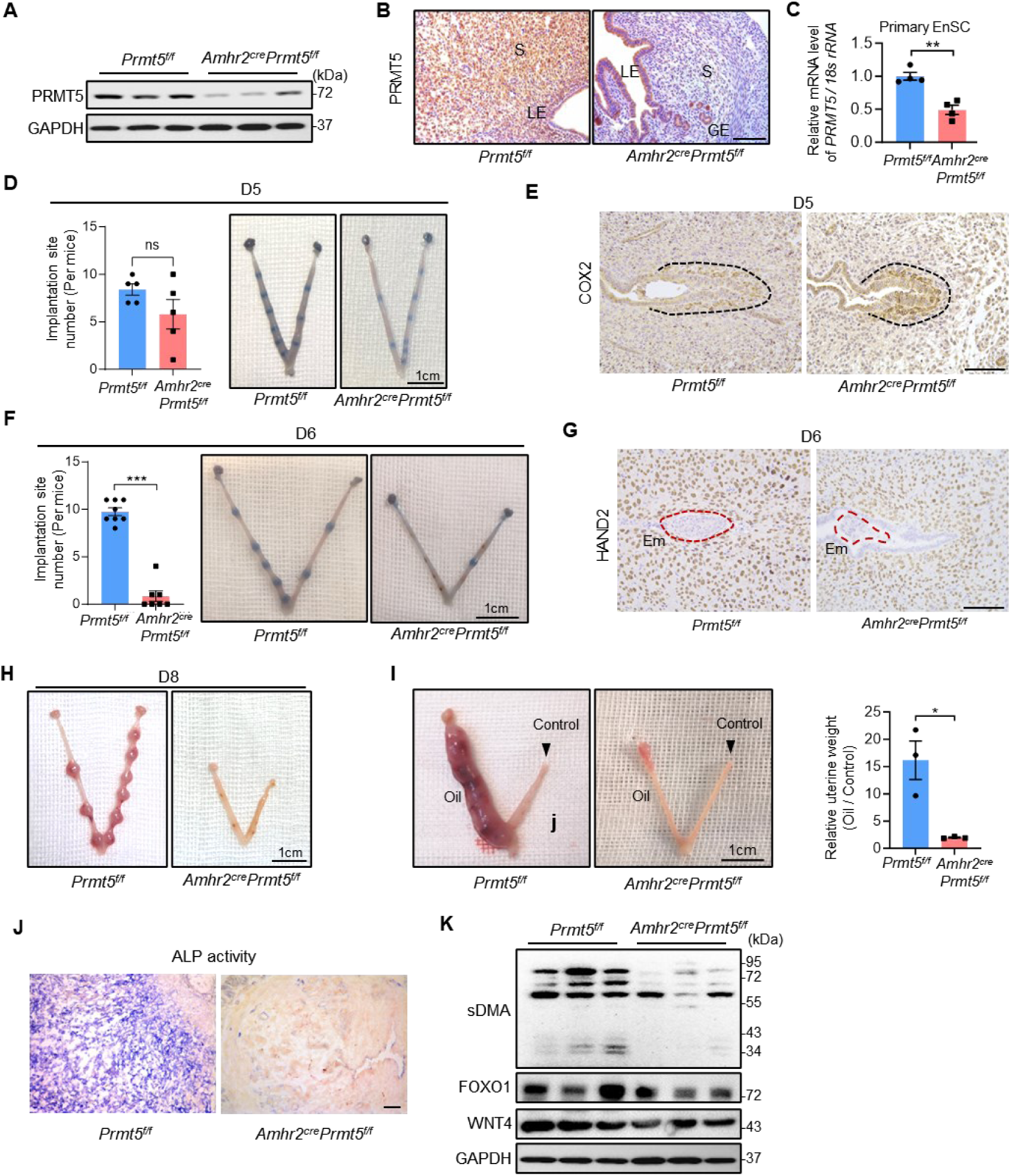
Loss of PRMT5 in endometrial stromal cells leads embryo implantation failure and decidualization defect in mice. (A) WB analysis of PRMT5 protein levels to reveal the knockout efficiency of *Prmt5^f/f^*and *Amhr2^cre^Prmt5^f/f^* uteri. (B) IHC staining of PRMT5 to reveal the knockout specificity of *Prmt5^f/f^* and *Amhr2^cre^Prmt5^f/f^* uteri. Scale bar, 100 μm. (C) qRT–PCR analysis of PRMT5 mRNA levels of primary EnSC of *Prmt5^f/f^* and *Amhr2^cre^Prmt5^f/f^* uteri to reveal the knockout efficiency. (D) Average number of implantation sites in *Prmt5^f/f^* and *Amhr2^cre^Prmt5^f/f^*mice on day 5 (D5) of pregnancy. Scale bar, 1 cm. (E) IHC staining of COX2 to show the embryo attachment reaction in implantation sites of *Prmt5^f/f^* and *Amhr2^cre^Prmt5^f/f^* mice on D5 of pregnancy. COX2 positive cells are circled by a black dotted line. Scale bar, 100 μm. (F) Average number of implantation sites in *Prmt5^f/f^*and *Amhr2^cre^Prmt5^f/f^* mice on D6 of pregnancy. Scale bar, 1 cm. (G) IHC staining of HAND2 to show the EnSC decidualization of *Prmt5^f/f^* and *Amhr2^cre^Prmt5^f/f^* mice on D6 of pregnancy. The embryo is circled by a red dotted line. Scale bar, 100 μm. (H) Representative images of D8 uteri from *Prmt5^f/f^* and *Amhr2^cre^Prmt5^f/f^*mice. Scale bar, 1 cm. (I) Gross morphology of unstimulated or oil-stimulated uterine side and the ratio of oil-stimulated to unstimulated uterine weight from *Prmt5^f/f^* and *Amhr2^cre^Prmt5^f/f^* mice. Scale bar, 1 cm. (J) Alkaline phosphatase (ALP) activity staining of oil-stimulated uterine side from artificial decidualization model of *Prmt5^f/f^*and *Amhr2^cre^Prmt5^f/f^* mice. Scale bar, 100 μm. (K) WB analysis of indicated protein levels to reveal impaired sDMA modification and decidualization regulators in oil-stimulated uterine side from artificial decidualization model of *Amhr2^cre^Prmt5^f/f^* mice. LE, luminal epithelium; GE, glandular epithelium; S, stroma; Em, embryo. Mean ± SEM. \**P* < 0.05, \*\**P* < 0.01, \*\*\**P* < 0.001, Student’s t test.

Both of *Prmt5^f/f^* and *Amhr2^cre^ Prmt5^f/f^* mice manifested normal embryo attachment reaction on the day 5 morning, which was visualized by blue dye reaction and COX2 positive cells around the blastocyst (Figure 1, D and E). *Prmt5^f/f^* mice exhibited a complete embryo implantation and obvious decidualization on day 6. However, the embryos in *Amhr2^cre^Prmt5^f/f^*mice did not invade through the luminal epithelium into the stroma on day 6, leading to implantation failure and arrested development, along with decidualization defect visualized by weakened HAND2 expression (Figure 1, F and G). Embryo growth and decidualization defect were further noted on day 8 (Figure 1H, and Supplemental Figure 1C). The potential ovarian defects were excluded by the fact of similar ovarian steroid hormones estradiol-17β (E2) and progesterone (P4), as well as normal follicle development in *Prmt5^f/f^*and *Amhr2^cre^ Prmt5^f/f^* mice (Supplemental Figure 2). In mice, decidualization was induced by embryo implantation in normal pregnancy or oil stimulation in artificially induced model(9, 26). To exclude the possible adverse effect of embryos, we induced artificial decidualization with intrauterine oil injection in the absence of the implanting embryo. *Amhr2^cre^ Prmt5^f/f^* mice exhibited a severe defect of decidualization as compared with *Prmt5^f/f^* mice, characterized by the decreased decidual tissue weight and Alkaline phosphatase (ALP) activity in the stroma (Figure 1, I and J). We also found that the overall symmetric dimethylarginine (sDMA) level and expression of key decidualization regulators (FOXO1 and WNT4) were significantly decreased in *Amhr2^cre^ Prmt5^f/f^* mice as compared with the *Prmt5^f/f^* mice (Figure 1K). Together, these results suggest that PRMT5 deficiency in EnSC is critically implicated in the embryo implantation failure and decidualization defect in the peri-implantation period.

### PRMT5 plays a crucial role in human endometrial stromal cell differentiation

We next determined the role of PRMT5 in human endometrial decidualization. In normal menstrual cycles, PRMT5 was expressed extensively in both epithelial and stromal cells. Epithelial and stromal PRMT5 levels were progressively increased in progesterone-dominant secretory phase compared with estrogen-dominant proliferative phase (n=24 for each group). We also observed the consistent increased contents of sDMA in secretory phase, indicated by a classic PRMT5 substrate Sm protein with molecular weight 26 kDa and various proteins with molecular weight more than 55 kDa (n=10 for each group) (27, 28) (Figure 2, A and B). The expression PRMT5 is upregulated, in a time-dependent manner, in primary human EnSC after treatment with medroxyprogesterone acetate (MPA) and 8Br-cAMP (Figure 2C). Further, knockdown of PRMT5 by adenovirus mediated shRNA in human EnSC resulted in suppressed mRNA expression levels of PRL and IGFBP1, two important decidual marker genes (Figure 2, D-F). The secreted PRL protein content in the supernatant was also decreased upon PRMT5 knockdown (Figure 2G). Morphological analysis with F-actin staining clearly showed that decidualized human EnSC displayed polygonal cell morphology, while PRMT5 knockdown resulted in more elongated fibroblast-like cells (Figure 2H). Together, these loss-of-function assays indicate that PRMT5 is indispensable for human endometrial decidualization.

**Figure 2.**
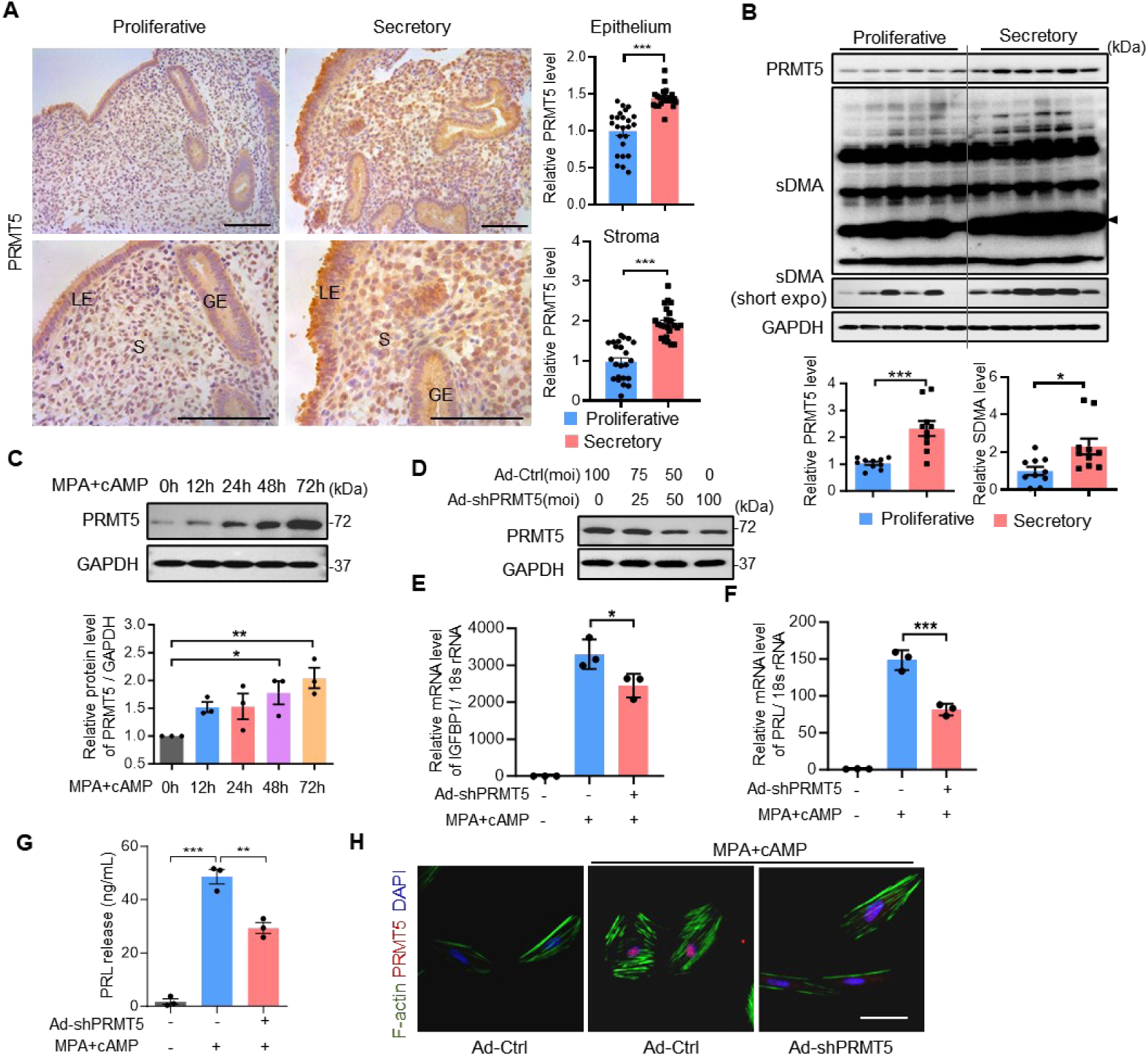
PRMT5 plays a crucial role in human endometrial stromal cell differentiation. (A) IHC staining of PRMT5 protein expression in proliferative endometrium (n = 24) and mid-secretory endometrium (n = 24) from normal fertile women. Scale bars, 100 μm. The integrated optical density (IOD) of each area from endometrium is analyzed using Image-Pro Plus 6.0. LE, luminal epithelium; GE, glandular epithelium; S, stroma. (B) WB analysis of PRMT5 and sDMA modified protein levels in proliferative endometrium (n = 6) and mid-secretory endometrium (n = 6) from normal fertile women. (C) WB analysis of PRMT5 protein level in human endometrial stromal cells (EnSC) treated with medroxyprogesterone acetate (MPA) and 8-bromo-cyclic adenosine monophosphate (8Br-cAMP; cAMP) for 0, 12, 24, 48 and 72 hours, respectively. (D) WB analysis of PRMT5 protein level in human EnSC infected with transfected with indicated multiplicity of infection (moi) of adenoviruses harboring shPRMT5 (Ad-shPRMT5). (E, F) qRT–PCR analysis of IGFBP1 and PRL mRNA levels of human EnSC transfected with Ad-shPRMT5 upon MPA and cAMP treatment for 3 days. (G) ELISA analysis of secreted PRL concentration in supernatant of human EnSC transfected with Ad-shPRMT5 upon MPA and cAMP treatment for 3 days. (h) IF staining of F-actin with phalloidin (green) and PRMT5 (red) in human EnSC transfected with Ad-shPRMT5 upon MPA and cAMP treatment for 3 days. Ad-Ctrl was used as the control virus. Scale bar, 50 μm. Mean ± SEM. \**P* < 0.05, \*\**P* < 0.01, \*\*\**P* < 0.001. Student’s t test in (A) and (B), ANOVA with Tukey’s multiple comparisons test in (C), (E) and (F).

### Both sDMA and PRMT5 are reduced in the endometrium of patients with RIF

The above data prompted us to investigate the potential role of arginine methylation in the pathology of endometrial decidualization. We first investigated the content of different arginine methylation in the endometrium from fertile women and patients with RIF. There was no significant difference in asymmetric dimethylarginine (aDMA) and monomethylarginine (MMA) contents between the two groups. However, levels of sDMA modification, as indicated by Sm protein with molecular weight 26 kDa and various proteins with molecular weight 34 kDa - 95 kDa, were markedly reduced in the RIF group compared with the fertile group (Figure 3A). Accordingly, in the two major type II PRMTs responsible for the generating sDMA, PRMT5 level was obviously decreased in the RIF group, but not PRMT9 (Figure 3B). The reduction of PRMT5 mRNA level was further supported by a greater sample size (n=22 for each group) (Figure 3C). Western blot data also confirmed the decreased level of endometrial PRMT5 in patients with RIF compared with the fertile women (n=24 for each group) (Figure 3D). Furthermore, the IHC assay revealed that reduced PRMT5 expression in RIF patients mainly occurred in the epithelia and stroma cells (n=24 for each group) (Figure 3E). The primary human EnSC isolated from endometrial tissues of RIF patients showed impaired decidualization after differentiation stimulus, as revealed by the mRNA levels of PRL and IGFBP1, and secreted PRL protein content. However, adenovirus-mediated overexpression of PRMT5 resulted in significant increase in the expression of PRL and IGFBP1 mRNA of EnSC from RIF patients by approximately 66% and 116%, respectively (Figure 3, F-H). Exogenous PRMT5 obviously promoted the content of secreted PRL protein in EnSC of RIF patients to 33.7 ng/mL (Control: 64.5 ng/mL, RIF: 21.4 ng/mL) and 59.0 ng/mL (Control: 103.2 ng/mL, RIF: 40.6 ng/mL), after 3 days and 6 days of differentiation stimulus, respectively (Figure 3I). Our results suggest that abnormally reduced levels of PRMT5 in the EnSC contribute to decreased sDMA contents and the decidualization defect in RIF patients.

**Figure 3.**
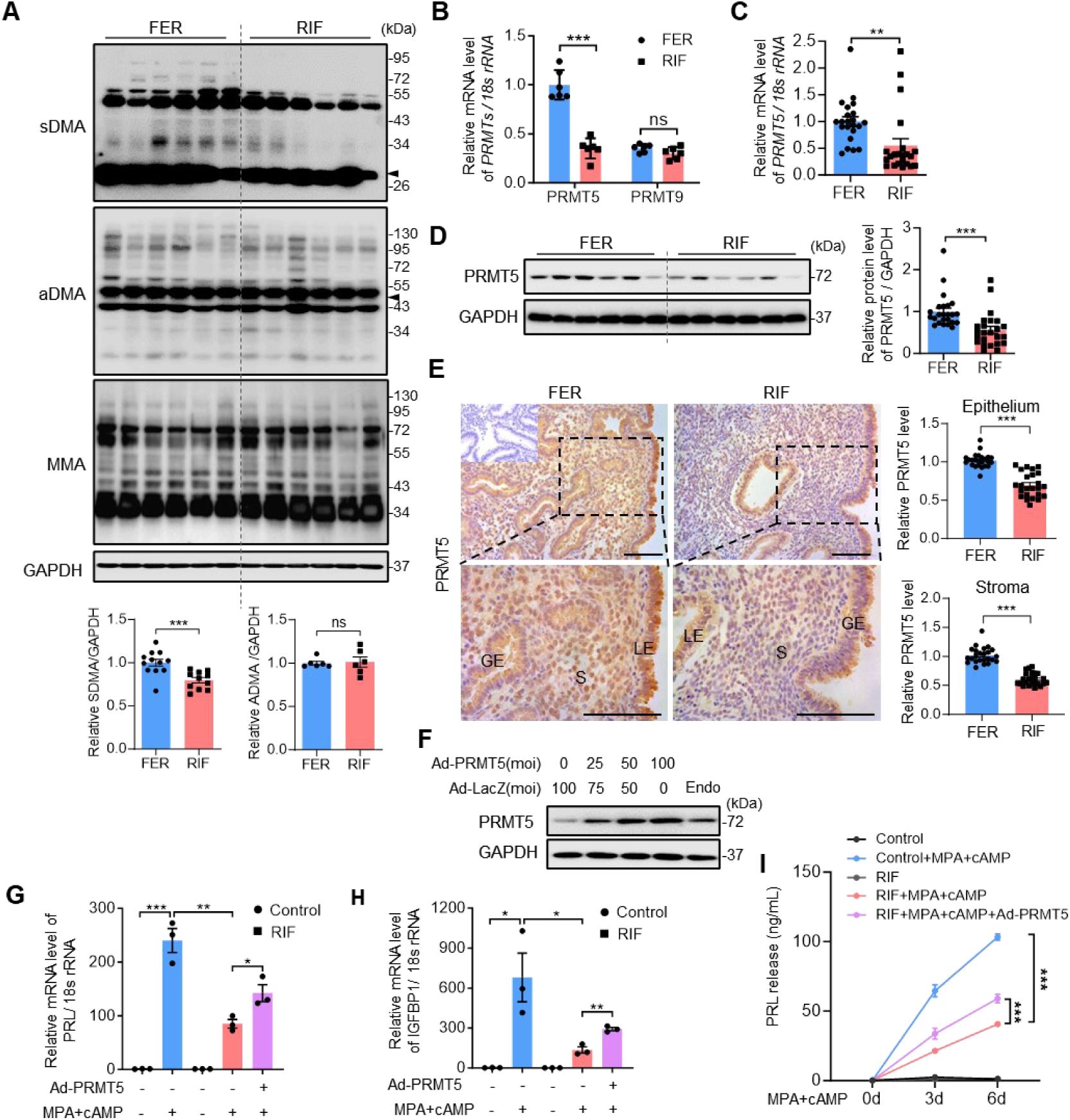
Both sDMA and PRMT5 are reduced in the endometrium of patients with RIF. (A) WB analysis of sDMA, aDMA and MMA modified protein levels and (B) qRT-PCR analysis of PRMT5 and PRMT9 mRNA levels in mid-secretory endometrium from infertile women with RIF (n = 6) and normal controls (n = 6). (C) qRT-PCR analysis of PRMT5 mRNA levels in mid-secretory endometrium from infertile women with RIF (n = 22) and normal controls (n = 22). (D) WB analysis and (E) immunohistochemistry staining of PRMT5 protein expression in mid-secretory endometrium from infertile women with RIF (n = 24) and normal controls (n = 24). (F) WB analysis of PRMT5 protein level in human EnSC infected with transfected with indicated moi of adenoviruses harboring PRMT5 (Ad-PRMT5). qRT–PCR analysis of (G) PRL and (H) IGFBP1 mRNA of human EnSC from infertile women with RIF and normal controls transfected with Ad-shPRMT5 upon MPA and cAMP treatment for 3 days. (I) ELISA analysis of secreted PRL concentration in supernatant of human EnSC from infertile women with RIF and normal controls transfected with Ad-shPRMT5 upon MPA and cAMP treatment for 3 days and 6 days. Ad-LacZ was used as the control virus. Scale bar, 100 μm. The integrated optical density (IOD) of each area from endometrium is analyzed using Image-Pro Plus 6.0. LE, luminal epithelium; GE, glandular epithelium; S, stroma. Mean ± SEM. \**P* < 0.05, \*\**P* < 0.01, \*\*\**P* < 0.001, Student’s t test. ANOVA with Tukey’s multiple comparisons test in (G) and (H). Two-way ANOVA with the Bonferroni multiple comparisons test in (J).

### PRMT5 regulates decidualization through targeting Nur77

To further understand the potential mechanism of action in stromal decidualization, we performed immunoprecipitation to identify PRMT5-interacting protein substrates using anti-PRMT5 antibody, followed by a mass spectrometry analysis in human EnSC. A total of 104 proteins were identified and these proteins include PRMT5 (bait), known PRMT5-interacting proteins such as Histone H2A and FAM120A(29, 30), and some novel interaction proteins, such as Histone H1.4, NR4A3, and Nur77 (also known as NR4A1) (Figure 4, A and B, and Supplemental Table 3). Further, RNA sequencing (RNA-Seq) was performed to compare the transcriptomes of control group (CTL), decidualization group (DEC), PRMT5 knockdown group (shPRMT5) and PRMT5 knockdown plus decidualization group (shPRMT5_DEC). The genes significantly downregulated after PRMT5 knockdown included aforementioned *IGFBP1* as well as other genes including *HBEGF*, *EZH2*, *MMP10*, *RBPJ*, *KLF12*, *DCN*, *BMP2*, *SOX4* and *RUNX2*, which are known to be critical for decidualization. The enrichment in extracellular matrix organization via Gene ontology (GO) enrichment analysis and epithelia mesenchymal transition pathway via Gene Set Enrichment Analysis (GSEA) indicated PRMT5 knockdown abolished the drastic differentiation procedure (Supplemental Figure 3, A-F). We found that 95 differentially expressed genes were common among the four groups, of which 38 were the target genes of Nur77 (*P*<0.001), which was found in PRMT5 immunoprecipitates (31) (Supplemental Figure 3, G and H). Co-immunoprecipitation assay confirmed the interaction of PRMT5 with Nur77 and other NR4A members in 293T cells (Figure 4, C and D). Albeit PRMT5 and Nur77 were universally localized in the cytoplasm and nucleus of EnSC, there was an obvious colocalization of PRMT5 and Nur77 in the nucleus, which was further enhanced after decidual differentiation (Figure 4, E and F). Knockdown of Nur77 blocked the promotion of PRMT5 on decidualization, indicating that Nur77 is a downstream effector of PRMT5 on decidualization (Figure 4, G and H). Notably, overexpression of Nur77 resulted in significant increase in the expression of PRL and IGFBP1 mRNA of PRMT5 knockdown EnSC by approximately 90% and 100%, respectively (Figure 4, I and J). Besides, exogenous Nur77 obviously promoted the content of secreted PRL protein in PRMT5 knockdown EnSC from 9.1 ng/mL to 20.47 ng/mL, and several key decidualization regulators expression (in which PRL, FOXO1 and HOXA10 are the known target genes of Nur77) (Figure 4K and Supplemental Figure 4). Together, our multi-omics evidence and loss and gain-of-function assays strongly suggest that Nur77 is a key mediator of PRMT5 in regulating EnSC decidualization.

**Figure 4.**
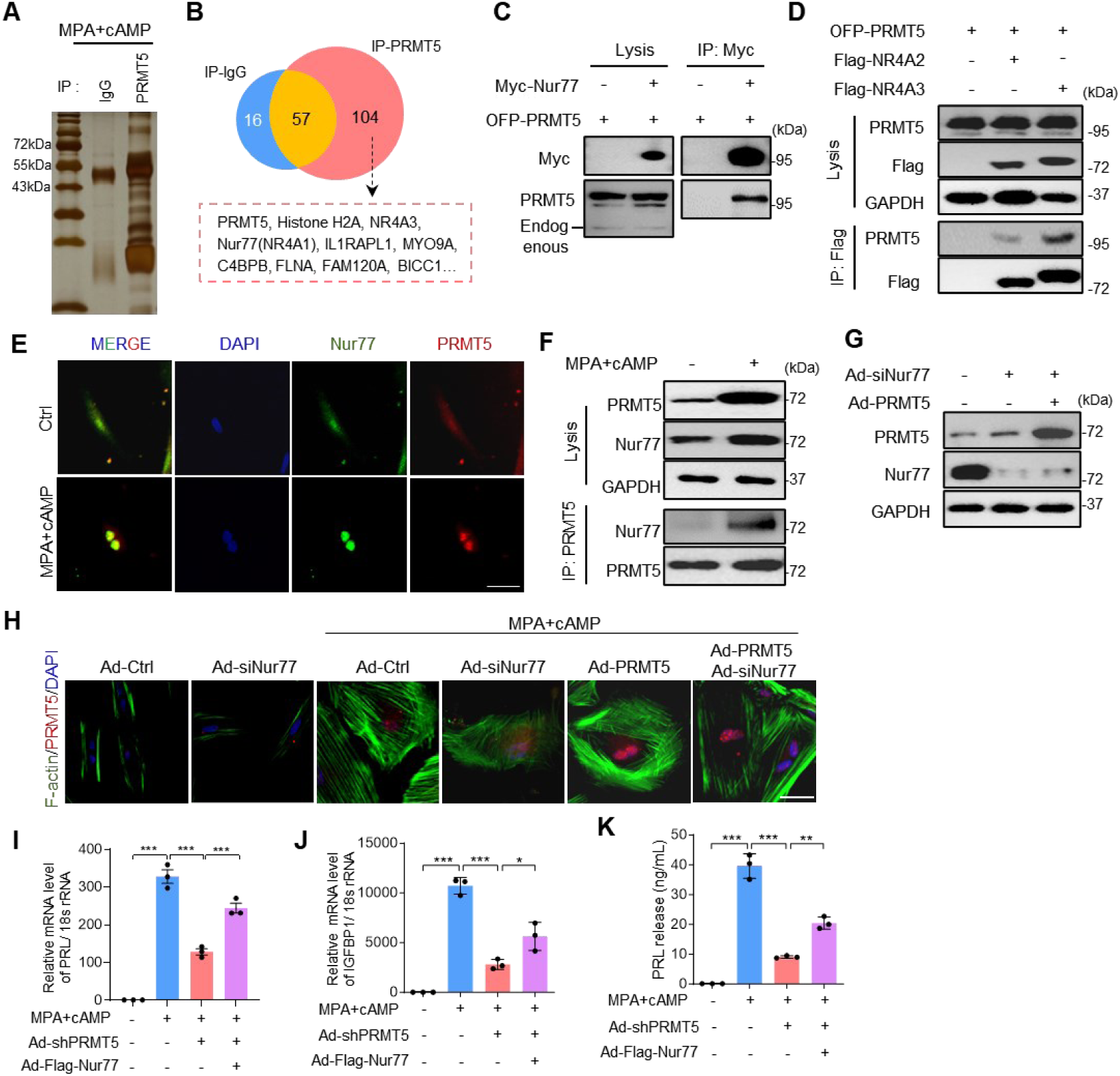
PRMT5 regulates decidualization through targeting Nur77. (A) Silver staining of proteins immunoprecipitated by IgG antibody and PRMT5 antibody using lysates of human EnSC treated with MPA and cAMP for 3 days. (B) Venn plot shows 104 proteins identified with confidence in the PRMT5 antibody group. (B) Co-IP and WB analysis of the interaction of exogenous OFP-PRMT5 with Flag-NR4A2 or Flag-NR4A3 in HEK293T cells. (D) Co-IP and WB analysis of the interaction of exogenous OFP-PRMT5 with Myc-Nur77 in HEK293T cells. (E) IF staining array for the localization of endogenous Nur77 (green) and PRMT5 (red) in human EnSC treated with MAP and cAMP. (F) Co-IP and WB analysis of the interaction of endogenous Nur77 and PRMT5 human EnSC treated with MAP and cAMP. (G) WB analysis of the expression level of Nur77 and PRMT5 in human EnSC transfected with Ad-siNur77 and Ad-PRMT5. (H) IF staining of F-actin with phalloidin (green) and PRMT5 (red) in human EnSC transfected with Ad-siNur77 and Ad-PRMT5 with MPA and cAMP treatment. (I, J) qRT–PCR analysis of PRL and IGFBP1 mRNA and (K) ELISA analysis of secreted PRL concentration of supernatant in human EnSC transfected with Ad-shPRMT5 and Ad-Flag-Nur77 with MPA and cAMP treatment. Ad-Ctrl and Ad-LacZ were used as the control virus. Scale bars, 50 μm. Mean ± SEM. \**P* < 0.05, \*\**P* < 0.01. ANOVA with Tukey’s multiple comparisons test.

### PRMT5 activates Nur77 through symmetric dimethylation of Nur77 at R346

To further investigate whether Nur77 is a bona fide substrate of PRMT5, we first mapped interaction domains of Nur77 with PRMT5. Nur77 consists of an N-terminal transactivation domain (NT), a middle DNA-binding domain (DBD), and a C-terminal ligand-binding domain (LBD). We found that PRMT5 bound to the NT and, to a lesser extent, the DBD and LBD domain (Figure 5A). On the other hand, overexpression of PRMT5 promoted Nur77-sDMA formation (Figure 5B). To investigate whether PRMT5-mediated methylation affects Nur77 transcriptional activity, we performed an electrophoretic mobility shift assay (EMSA). As shown in Figure 5C, PRMT5 knockdown decreased the binding of Nur77 to nerve growth factor-induced B factor response element (NBRE) oligonucleotides. Luciferase reporter assay showed that PRMT5 knockdown attenuated Nur77 dependent NBRE activation in human EnSC (Figure 5D). Further, primary mouse EnSC isolated from *Amhr2^cre^Prmt5^f/f^*mice exhibited weaker activity of NBRE-driven luciferase than that from *Prmt5^f/f^*mice (Figure 5E). PRMT5 deficiency led to an increased degradation of Nur77 protein (Figure 5F). In addition, we found that PRMT5 deficiency disturbed the nuclear location of Nur77 in decidual EnSC (Figure 5G). To identify PRMT5 methylation sites in Nur77, we analyzed the protein sequence of Nur77 using methylation prediction tools including PRmePred and GPS-MSP. Four arginine residues ranked top scores were selected for further analyses, but only the substitution of arginine (R) to lysine (K) at 346 (346K) completely blocked Nur77-sDMA formation (Figure 5H). Importantly, in vitro methylation assays demonstrated that PRMT5 directly methylates Nur77 DBD, but not NT, LBD or DBD-346K mutant (Figure 5I). R346 and its surrounding amino acid sequences exhibited remarkable conservation among diverse species, indicating potential evolutionary significance (Figure 5J). Nur77 exhibited both higher NBRE luciferase activity and protein stability compared with R346K mutant (Figure 5, K and L). These results suggest that PRMT5 catalyzes symmetric dimethylation of Nur77 at R346 and increases the transcriptional activity of Nur77 in EnSC.

**Figure 5.**
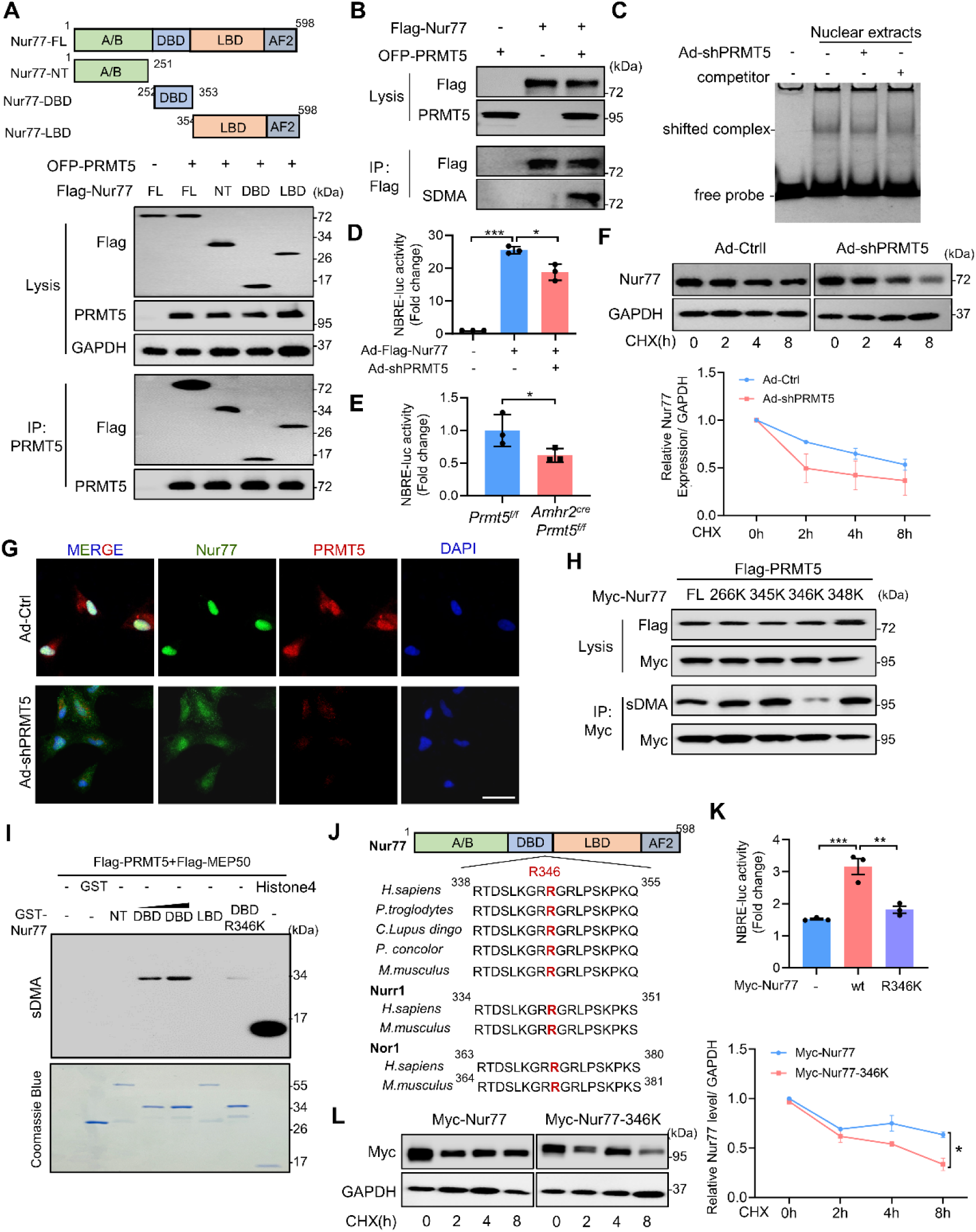
PRMT5 activates Nur77 through symmetric dimethylation of Nur77 at R346. (A) Schematic representation of Flag-tagged human Nur77 truncation derivatives, including NT, DBD and LBD. Co-IP and WB analysis of the interaction between full-length PRMT5 and Nur77 fragments in HEK293T cell. Three domains of Nur77 interact with PRMT5. (B) Exogenous Myc-Nur77 was immunoprecipitated from HEK293T cell transfected OFP-PRMT5 plasmid by Myc antibody, and the arginine methylation status of Nur77 was examined with a sDMA antibody. (C) Electrophoretic Mobility Shift Assay of binding capacity of extracts from human EnSC transfected with Ad-shPRMT5 to cy5 labeled NBRE DNA probe, unlabeled NBRE DNA probe was used as competitor. (D) Human EnSCs transfected with Ad-Flag-Nur77 and Ad-shPRMT5 and (E) EnSCs from *Prmt5^f/f^*and *Amhr2^cre^Prmt5^f/f^* mice were subjected to the detection of NBRE-luciferase activity. (F) Human EnSCs transfected with Ad-shPRMT5 were treated with 50 μg/mL cycloheximide (CHX) for 2, 4, and 8 hours. The cell extracts were subjected to Western blotting. The level of the remaining Nur77 was normalized to that of GAPDH and plotted relative to the level at the 0-hour time point. (G) IF staining of Nur77 and PRMT5 shows the intracellular localization of Nur77 in human EnSCs transfected with Ad-shPRMT5. (H) WB analysis of immunoprecipitated proteins, obtained from 293T cells transiently transfected with plasmids expressing Myc tagged Nur77, Nur77 mutants including R266K, R345K, R346K, and R348K. (I) Purified GST-Nur77 truncation derivatives, including NT, DBD, LBD and DBD R346K mutant, Flag-PRMT5 and Flag-MEP50 fusion protein were subjected to in vitro methylation analysis with a sDMA antibody to detect Nur77 methylation. Histone4 protein was used as the positive control. (J) Alignment of the consensus Nur77 amino acid sequences around the arginine 346 residue highlighted in red among various species. (K) NBRE-luciferase activity analysis of HEK293T exogenously expressing Myc-Nur77 and Myc-Nur77-R346K. (L) HEK293T exogenously expressing Myc-Nur77 and Myc-Nur77-R346K were treated with CHX for 2, 4, and 8 hours, then subjected to Western blotting. The level of the remaining Myc tagged protein was normalized to that of GAPDH and plotted relative to the level at the 0-hour time point. Mean ± SEM. \**P* < 0.05, \*\**P* < 0.01, \*\*\**P* < 0.001. Student’s t test in E. ANOVA with Tukey’s multiple comparisons test in (D) and (K). Two-way ANOVA with the Bonferroni multiple comparisons test in (F) and (L).

### PRMT5-mediated Nur77 methylation impacts phosphorylation of Nur77 by AKT

Phosphorylation of NR4A by AKT has been shown to decrease the transcriptional activity of Nur77 (32). Thus, we speculated that there may be a crosstalk between the methylation and phosphorylation of Nur77. Kyoto Encyclopedia of Genes and Genomes (KEGG) analysis and GSEA revealed an enrichment of PI3K/AKT pathway in PRMT5 knockdown decidualized EnSC (Figure 6, A and B). The increased level of phosphorylated Nur77 at Serine 351 (S351) and phosphorylated AKT confirmed that knockdown of PRMT5 in EnSC led to AKT activation and Nur77 phosphorylation, along with decreased levels of IGFBP1, HOXA10 and FOXO1 (Figure 6C). Furthermore, *Amhr2^cre^ Prmt5^f/f^* mice showed increased Ki67 positive EnSCs and phosphorylation levels of AKT and Nur77 in the uterus compared to the control (Figure 6, D and E). We performed an immunoprecipitation assay in EnSC isolated from *Prmt5^f/f^*mice and *Amhr2^cre^Prmt5^f/f^* mice, and observed that PRMT5 knockout attenuated Nur77 methylation but elevated Nur77 phosphorylation (Figure 6F). Besides, methylation-deficient Nur77 mutant 346K exhibited a large rise in the AKT mediated phosphorylation of Nur77, while the interaction between Nur77 and PRMT5 was unaffected obviously in the methylation-deficient Nur77 346K mutant (Figure 6, G and H). Furthermore, we demonstrated that the binding of Nur77 to pAKT was enhanced in PRMT5 knockdown EnSC (Figure 6I), and our bioinformatics analysis illustrated that Nur77 formed multiple hydrogen bond interactions with PRMT5, including the amino acids around R346, whereas Nur77 had fewer interactions with AKT (Supplemental Figure 5), indicating that PRMT5 mediated R346 methylation of Nur77 affects S351 phosphorylation level through changing the interaction between pAKT and Nur77. Taken together, these data indicate that there exists a crosstalk between PRMT5-mediated Nur77 R346 methylation and AKT-mediated Nur77 S351 phosphorylation in EnSC.

**Figure 6.**
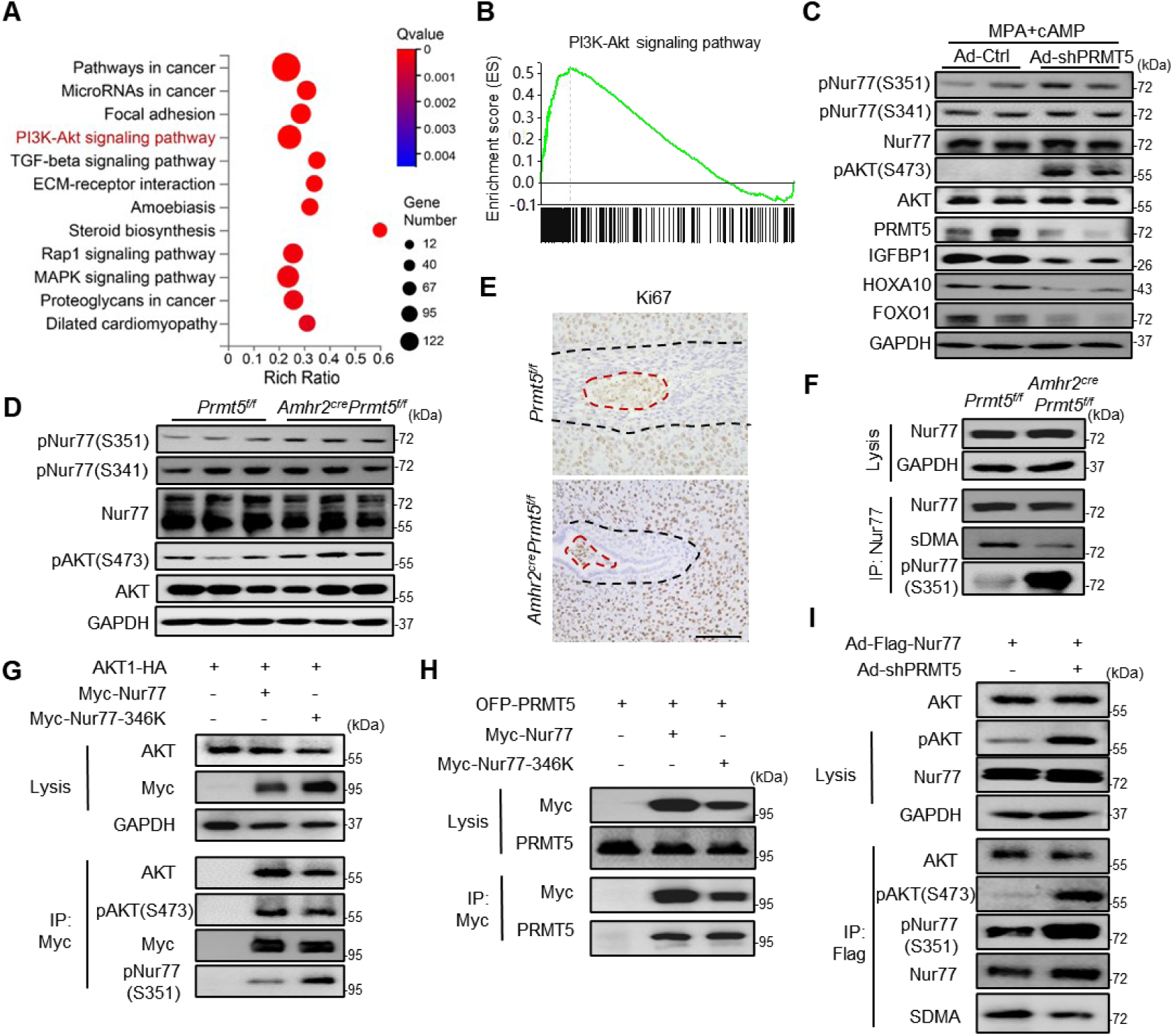
PRMT5-mediated Nur77 methylation impacts phosphorylation of Nur77 by AKT. (A) KEGG enrichment pathway analysis of differential expression genes between control and PRMT5 knockdown human EnSC treated with MPA and cAMP. (B) The “PI3K-AKT signaling” gene set was enriched in PRMT5 knockdown decidualized EnSC group according to GSEA. WB analysis of phosphorylated AKT Ser473 (pAKT S473), pNur77 S341 and pNur77 S351 in (C) Ad-shPRMT5 transfected human EnSC with treatment of MPA and 8Br-cAMP and (D) *Amhr2^cre^Prmt5^f/f^* mice uteri. (E) IHC analysis of Ki67 positive EnSC in *Prmt5^f/f^* and *Amhr2^cre^Prmt5^f/f^*mice uteri on day 6 of pregnancy. Ki67 positive cells are circled by a black dotted line, and the embryo is circled by a red dotted line. Scale bar, 100 μm. (F) IP and WB assay for *Amhr2^cre^Prmt5^f/f^* mice uteri to reveal the relationship of Nur77-sDMA and pNur77 S351. (G) Co-IP and WB assay for HEK293T transfected with AKT1-HA and Myc-Nur77 or Myc-Nur77-346K to reveal the role of R346 mutant on the interaction between Nur77 and AKT or pAKT(S473). (H) Co-IP and WB assay for HEK293T transfected with OFP-PRMT5 and Myc-Nur77 or Myc-Nur77-346K to reveal the role of R346 mutant on the interaction between Nur77 and PRMT5. (I) Co-IP and WB assay for human EnSC transfected with Ad-Flag-Nur77 and Ad-shPRMT5 to reveal the role of PRMT5 knockdown on the interaction between Nur77 and AKT or pAKT(S473).

### PRMT5 deficiency leads to aberrant proliferation of EnSC

Due to enhanced AKT signaling activity and Ki67 positive cells after PRMT5 knockdown, we next determined the potential role of PRMT5 in modulating the balance between methylation and phosphorylation of Nur77 in the transition from proliferation to differentiation of EnSC. Primary human EnSCs were cultured in 10%FBS, 2.5%FBS or 2.5%FBS with MPA and 8Br-cAMP. 10% FBS culture represented the proliferation stage, which contributed to the lowest level of PRMT5 and Nur77-sDMA, but the highest level of pAKT, pNur77 and the percentage of Ki67 positive EnSC. Instead, 2.5%FBS with MPA and 8Br-cAMP represented the differentiation stage, which contributed to the highest level of PRMT5 and Nur77-sDMA, but the lowest level of pAKT, pNur77 and the percentage of Ki67 positive EnSC (Figure 7, A and B). In addition, knockdown of PRMT5 contributed to amplified sDMA modification in the nucleus, elevated pNur77 level in the cytoplasm, and promoted proliferation of EnSC (Supplemental Figure 6). The negative relationship between Ki67 positive percentage with PRMT5 expression in EnSC was observed in the endometrial samples obtained on different days after the luteinizing hormone surge (LH+3, LH+5, and LH+9) (Figure 7C). Importantly, all PRMT5 positive EnSCs are Ki67 negative *in vitro* and *in vivo* (Figure 7, B and D). Furthermore, the levels of PRMT5 and Nur77-sDMA were increased during LH+3 to LH+9, while the levels of pAKT and pNur77 were decreased along with the shift from proliferation to differentiation of EnSC (Figure 7E). We next determined the relationship between PRMT5, pAKT, and pNur77 in the endometrial samples of RIF patients. IHC assay of adjacent slices showed the decreased PRMT5 expression and increased pNur77 levels in the same stromal cells (Figure 7F). Western blot data further supported that the endometrium samples from RIF with deficient PRMT5 manifested decreased Nur77-sDMA level, but increased pNur77 level (Figure 7G). These results suggest that deficient PRMT5 expression contributes to the aberrant proliferation of EnSC, which exhibits an abnormal elevation of phosphorylated Nur77 in RIF patients.

**Figure 7.**
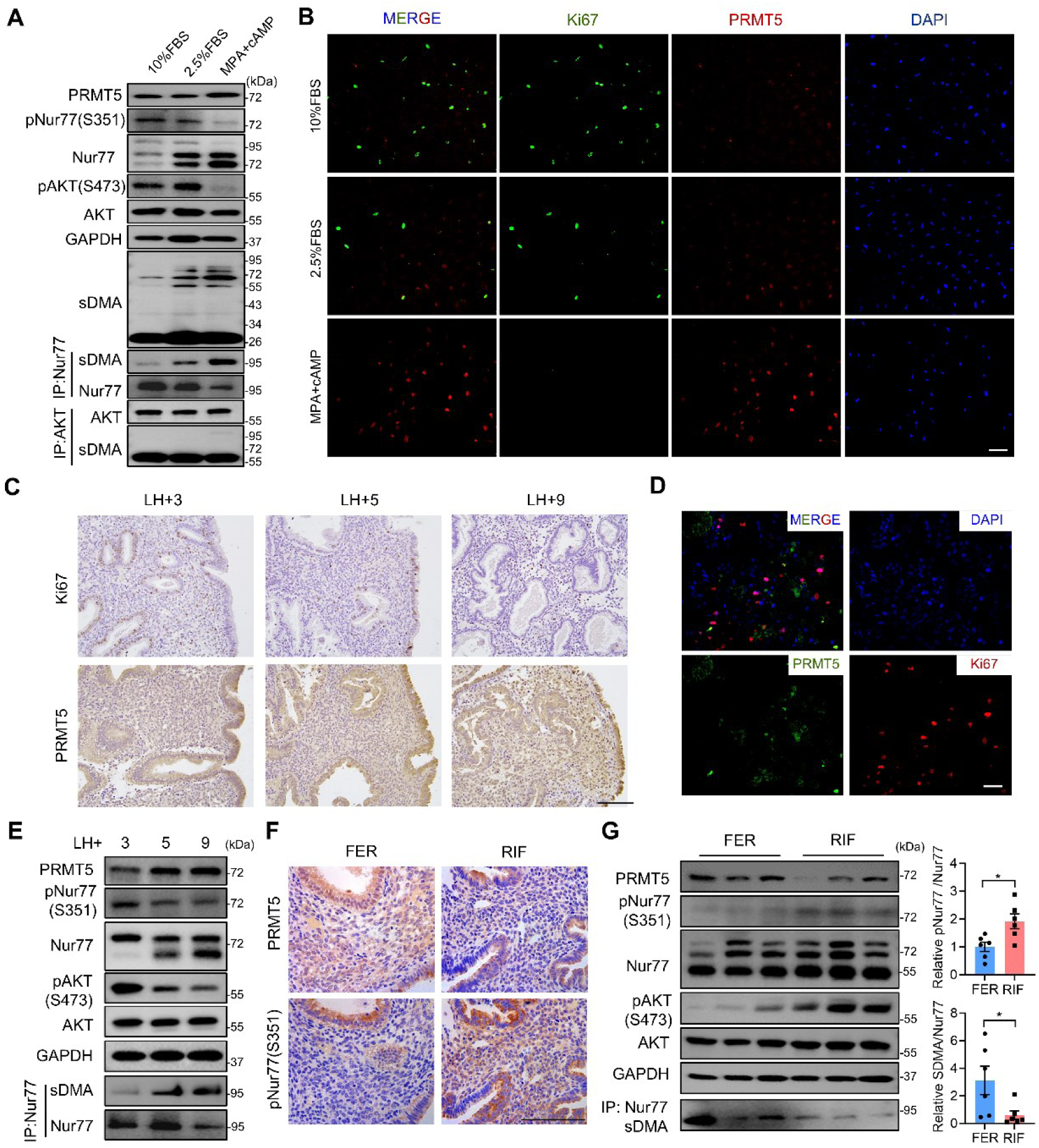
PRMT5 deficiency leads to aberrant proliferation of EnSC. (A) WB analysis of PRMT5, pNur77(S351), pAKT(S473) and sDMA modified Nur77 and (B) IF staining of Ki67 and PRMT5 in the human EnSC cultured in 10% FBS, 2.5% FBS and 2.5% FBS with MAP+8Br-cAMP. (C) IHC staining of Ki67 and PRMT5 in human endometrium from fertile women sampled in LH+3, LH+5 and LH+9. (D) IF staining of Ki67 and PRMT5 in the human endometrium from fertile women. (E) WB analysis of PRMT5, pNur77(S351), pAKT(S473) and sDMA modified Nur77 in human endometrium from fertile women sampled in LH+3, LH+5 and LH+9. (F) IHC analysis of mid-secretory endometrial PRMT5 and pNur77(S351) protein expression in women with RIF versus normal controls. (G) WB analysis of PRMT5, pNur77(S351), pAKT(S473) and Nur77-sDMA protein levels in mid-secretory endometrium from infertile women with RIF and normal controls. Scale bars, 100 μm.

### Nur77 methylation and phosphorylation modulates the decidualization

Finally, we explored the functional significance of Nur77 phosphorylation and methylation in decidual regulation. We first demonstrated that exogenous expression of Nur77 could partly rescue impaired decidualization of the primary EnSCs isolated from patients with RIF (Figure 8, A and B). After treating human EnSCs with Nur77 pharmacological activator Cytosporone B (CsnB) (33), we observed an improvement on the decidualization of primary human EnSC from RIF patients (Figure 8, C-E). Furthermore, inhibition of Nur77 phosphorylation with AKT pharmacological inhibitor MK2206 led to an inhibition of pAKT and pNur77 levels (Figure 8F). Importantly, MK-2206 treatment promoted Nur77 transcription activity, along with a mild increase of Nur77 methylation (Figure 8, G and H). We further demonstrated that MK2206 obviously rescued the decidualization defect of PRMT5 knockdown human EnSCs to the similar extent as CsnB (Figure 8, I and J). However, methylation deficient Nur77 mutant with elevated phosphorylation barely improves IGFBP1 and PRL expression in PRMT5 knockdown human EnSCs (Supplemental Figure 7). We speculated that a peptide with R351 (named as Pep1) could weaken AKT-mediated Nur77 phosphorylation and function as the competitive substrate, while a peptide with R346K mutant and R351 (named as Pep2-mut) could preserve the endogenous methylation of Nur77 (Figure 8K). Our data showed that Pep1 and Pep2-mut promoted the Nur77 transcriptional activity (Figure 8L). More importantly and interestingly, Pep2-mut exhibited an obvious promoting effect on IGFBP1 and PRL mRNA expression during the differentiation of EnSCs, which was superior to the Pep1 (Figure 8, M and N). Mechanistically, we did observe that Pep2-mut treatment contributed to decreased endogenous pNur77, while did not disturb pAKT or Nur77-sDMA obviously (Figure 8O). Together, our data indicate the significance of the methylation and phosphorylation on Nur77-mediated decidualization and highlight the PRMT5/AKT/Nur77 axis as promising therapeutic targets to modulate the decidualization.

**Figure 8.**
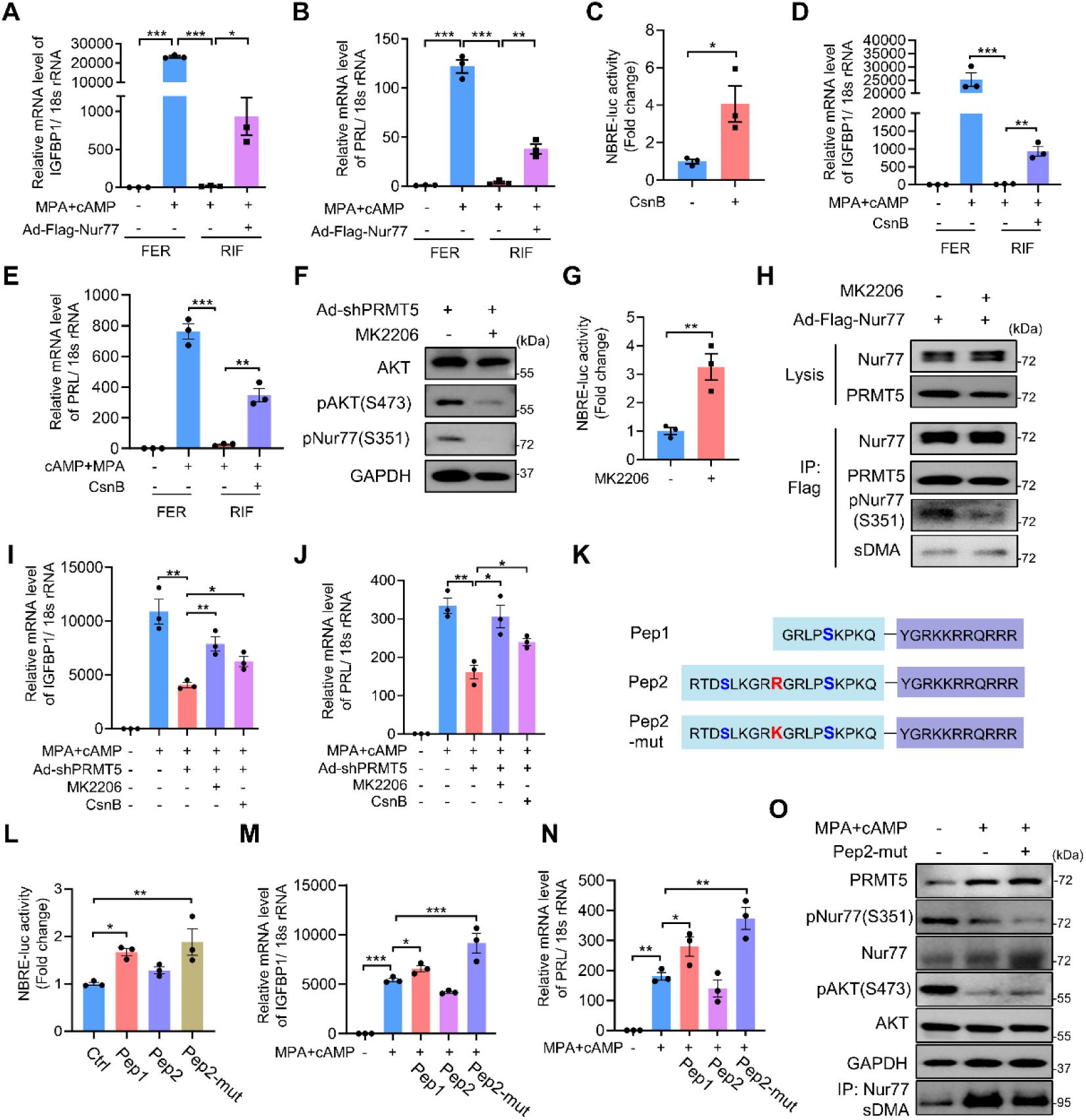
Impaired phosphorylation of Nur77 in the endometrium of women with unexplained RIF. qRT-PCR analysis of (A) IGFBP1 and (B) PRL mRNA level to reveal the role of exogenous overexpressing Nur77 on the differentiation capacity of human EnSC from RIF patients. (C) NBRE-luciferase activity analysis of human EnSC upon Cytosporone B (CsnB) treatment. qRT-PCR analysis of (D) IGFBP1 and (E) PRL mRNA level to reveal the role of CsnB on the differentiation capacity of human EnSC from RIF patients. (F) WB assay for pAKT(S374) and pNur77 (S351) in human EnSC transfected with Ad-shPRMT5 upon MK2206 treatment. (G) NBRE-luciferase activity analysis of human EnSC upon MK2206 treatment. (H) Co-IP and WB for human EnSC overexpressing Flag-Nur77 with MK2206 treatment to reveal the role of MK2206 on the interaction between Nur77 and PRMT5, and regulation of Nur77-sDMA and pNur77 S351. qRT-PCR analysis of (I) IGFBP1 and (J) PRL mRNA level to reveal the role of MK2206 or CsnB on the differentiation capacity of human EnSC with PRMT5 knockdown. (K) Schematic representation of Nur77-derived peptides including Pep1, Pep2 and Pep2-mut. (L) NBRE-luciferase activity analysis of human EnSC upon Nur77-derived peptides treatment. RT-PCR analysis of (M) IGFBP1 and (N) PRL mRNA level to reveal the role of Nur77-derived peptides on the differentiation capacity of human EnSC with MPA and 8Br-cAMP treatment. (O) WB analysis of PRMT5, pNur77(S351), pAKT(S473) and Nur77-sDMA protein levels in human EnSC treated with Nur77-derived peptide Pep2-mut. Mean ± SEM. \**P* < 0.05, \*\**P* < 0.01, \*\*\**P* < 0.001. Student’s t test in C and G. ANOVA with Tukey’s multiple comparisons test in (A, B, D, E, I, J, L, M, and N).

## Discussion

PRMTs are highly conserved from yeast to human and are responsible for arginine methylation by transferring methyl groups from S-adenosylmethionine to a guanidine nitrogen of arginine in proteins, which can be classified into three types according to their catalytic activity: type I (PRMT1, PRMT2, PRMT3, PRMT4, PRMT6, and PRMT8) and type II (PRMT5 and PRMT9) enzymes carry out the formation of MMA (by type III enzyme PRMT7) as an intermediate before the establishment of aDMA or sDMA, respectively(34). A recent study showed that aDMA content and PRMT3 expression were increased in the decidua of recurrent miscarriage patients, primarily in macrophages but not in stromal cells of the decidua(35). However, the role of arginine methylation with corresponding PRMT in the endometrium of RIF patients has not been fully uncovered. In this study, by screening these three types of methylation in the endometrial samples, we found that sDMA is significantly reduced in the RIF patients and that PRMT5 is a major contributor involved in this process. Furthermore, we provided genetic evidence that PRMT5 governs the EnSC decidualization to guarantee normal embryo implantation. Mechanically, PRMT5 methylates R346 residue to promote the transcriptional activity of Nur77, which is a key transcriptional factor involved in decidualization. More importantly and interestingly, we revealed a crosstalk between PRMT5-mediated Nur77 methylation at R346 and AKT-mediated Nur77 phosphorylation at S351, and provided the proof of concept that modulating the dynamic balance between the methylation and phosphorylation of Nur77 is a promising target therapy for embryo implantation with EnSC decidualization defect.

PRMT5 has been essentially implicated in cancer cell survival, proliferation, migration and metabolism, while the roles of PRMT5 in non-tumor studies, including immune response, ovarian follicle development, angiogenesis, have begun to be explored(36–39). To clarify the potential role of PRMT5 in the endometrium, we utilized PRMT5 knockdown human EnSCs and stromal conditional PRMT5 deletion mouse model. Defective decidualization of endometrial stromal cells *in vitro* was determined by aberrant morphological change and decreased levels of IGFBP1 and PRL in the PRMT5 knockdown EnSC. Defective decidualization *in vivo* was obviously observed in PRMT5 stromal conditional knockout mice. Normal embryo attachment reaction occurs in the PRMT5 conditional KO mice on day 5, but the embryos were unable to invade through the luminal epithelium into the stroma on day 6, leading to embryo implantation failure, which is similar to the clinical outcome of the RIF patients. Decidual defects have been revealed by our and other groups to be associated with RIF in recent years(10–14). We have indeed observed that the EnSCs in PRMT5 KO mice presented a decidual defect, with a smaller size and lower expression levels of HAND2. Instead of cyclic decidualization in human, mouse decidualization was induced by normal embryo implantation or artificial oil stimulation(9, 26). Utilizing an artificial decidualization model to exclude the embryo factor, we further demonstrate that stromal PRMT5 deficiency is mainly responsible for the decidualization defect. Our previous studies utilizing GSK591, a specific inhibitor of PRMT5, demonstrated that PRMT5 is essentially involved in the regulation of human EnSC decidualization(24). Here, we provided the genetic evidence for the indispensable function of PRMT5 in guaranteeing normal embryo implantation by governing EnSC decidualization.

PRMT5 is an epigenetic modifier that plays a key role in transcriptional regulation by catalyzing symmetric dimethylarginine of histone proteins, including H2AR3, H3R8, and H4R3(40). Besides to histones and chromatin remodeling complexes, PRMT5 regulates gene expression through methylating non-histone proteins such as transcription factors and signaling molecules, including p53, androgen receptor (AR), epidermal growth factor receptor (EGFR), and p65 subunit of NF-κB(41–44). Herein, we identified the potential interacting substrates of PRMT5 using immunoprecipitation followed by mass spectrometry analysis. We identified Nur77 as an ideal target when combining with the transcriptomic bioanalysis of PRMT5 knockdown EnSCs. Nur77 (also known as NR4A1 or TR3) is an orphan nuclear receptor that belongs to the steroid/thyroid/retinoid receptor superfamily. Recently, we found that that Nur77 is a key transcriptional factor involved in embryo implantation during the peri-implantation period, by transcriptionally promoting several key regulators, including β3-integrin, HOXA10, FOXO1 and PRL, to modulate the endometrial receptivity and stromal decidualization (31, 45–47). Here, by performing both loss and gain-of-function assays, we showed that Nur77 is a key mediator of PRMT5 to regulate EnSC decidualization. The transcriptional activity of Nur77 is regulated by phosphorylation, which is mainly mediated by AKT at Ser 351(32). Activation of PI3K/AKT pathway has been proven to compromise human EnSC decidualization(48). Although several phosphorylation sites of Nur77 were identified(45, 49), the functional significance of other PTMs such as methylation in human EnSC decidualization is unknown. In this study, we demonstrated that PRMT5 catalyzes symmetric dimethylation of Nur77 at R346, which is close to S351 (Nur77 346-351: RGRLPS). Previous studies have shown that PRMT1 mediated methylation of FOXO1 and BCL2 within an Akt consensus phosphorylation motif (RxRxxS/T) inhibits AKT mediated phosphorylation(21, 50). An intriguing finding in our study is that Nur77 is capable of being fine-tuned by both methyltransferase PRMT5 and phosphokinase AKT during the transition from proliferation to differentiation of EnSCs. Numerous studies have found the interplay between protein methylation and phosphorylation, which modulate the function regulation of target proteins. For instance, PRMT5-mediated methylation on R1175 of EGFR positively modulates EGF-induced EGFR trans-autophosphorylation at Tyr 1173, hence suppressing EGFR-mediated ERK activation(44). PRMT5-mediated methylation of Sterol regulatory element-binding protein 1a (SREBP1a) at R321 prevents phosphorylation of SREBP1a on Ser 430 by GSK3β, thereby promoting transcriptional activity(51). MAPK1 kinase mediated Lymphoid-specific helicase (LSH) at S503 antagonizes LSH R309 methylation by PRMT5, which eventually promotes stem-like properties in lung cancer(52). PRMT5 mediated methylation suppresses MST2 autophosphorylation and kinase activity by blocking its homodimerization, thereby inactivating Hippo signaling pathway in pancreatic cancer(53). Herein, our data demonstrated that methylation of Nur77 weakens phosphorylation, which is partly through changing the interaction between AKT and Nur77. Our three-dimensional structure prediction of PRMT5/Nur77/AKT complex supports that Nur77 tends to form more hydrogen bond interactions with PRMT5. In actuality the mutation R346 alone promotes AKT mediated phosphorylation of Nur77, but not obviously affect the interaction between AKT and Nur77. However, PRMT5 deficiency leads to both of increased Nur77 phosphorylation and enhanced interaction between AKT and Nur77. This may be explained by the fact that PRMT5 bounds to the NT and, to a lesser extent, the DBD and LBD domain of Nur77, hence the mutation R346 alone is not able to abolish the interaction between Nur77 and PRMT5. Although the detailed mechanism of AKT signaling activation in PRMT5 deficient EnSC has not been clarified here, a recent study reported that PRMT5 promotes AKT activation by catalyzing symmetric dimethylation of AKT1 at R391 in MCF7 cells(54). However, we did not observe the expression of AKT-sDMA in proliferative or differentiated EnSC (Figure 7A). In addition, it is yet unclear whether Nur77 is subjected to undergo other post-translational modifications, such as ubiquitination or acetylation, and whether other PTMs crosstalk with Nur77 methylation or phosphorylation, eventually affecting Nur77 transcriptional activity. Nevertheless, the specific mechanisms still need to be further investigated to further clarify the complex regulation network among PRMT5, AKT and Nur77 (or its homologous protein Nurr1 and Nor1).

The human endometrium undergoes a complex series of organized proliferative and secretory changes to prepare for embryo implantation(55). EnSC exits from cell cycle in response to the decidual differentiation signals, which is mediated by several key transcription factors, including CEBPB, FOXO1, STAT5 and ATF3. These transcription factors, in turn, are responsible for the expression of decidual marker genes PRL and IGFBP1(11, 56, 57). Our previous studies have found that Nur77 transcriptionally regulates FOXO1 and PRL(31, 47). Herein, we reported a transition from phosphorylation to methylation of Nur77 in EnSCs during the decidual transformation period, and EnSCs of RIF patients and PRMT5 deletion mice exhibit impaired decidualization and hyperproliferation. However, a recent study reports that PRMT5 knockout inhibits proliferation and promotes premature differentiation of embryonic myoblasts by promoting FOXO1 cytoplasmic accumulation, indicating the specific role of PRMT5 in various cells and organs(58). Based on the above studies, we postulated that modulating the balance of Nur77 methylation and phosphorylation to promote its transcriptional activity may impose a positive effect on the impaired decidualization. Pharmacological inhibition of AKT activation with MK2206 suppresses Nur77 phosphorylation, while promoting Nur77 methylation and its transcriptional activity. MK2206 rescues the decidualization defect in PRMT5 knockdown human EnSCs, which is not inferior to the pharmacological activation of Nur77 by CsnB. Importantly, we synthesized a Nur77-derived peptide with R346 mutant and R351 (Pep2-mut) which promotes the Nur77 transcriptional activity and EnSC decidualization. Our dada indicates that R351 in Pep2-mut could disturb the interaction of Nur77 with AKT as the competitive substrate and weaken endogenous Nur77 phosphorylation, but R346 mutant in Pep2-mut does not damage the endogenous Nur77 methylation by PRMT5 obviously. Pep2-mut may tip the balance of these two modifications toward methylation, which can efficiently improve the decidualization, yet the detailed mechanism needs to be further clarified in the future.

Taken together, our study provides the genetic evidence for the indispensable function of PRMT5 in guaranteeing endometrial stromal cell decidualization and embryo implantation by modulating the crosstalk between methylation and phosphorylation of Nur77. We put forward a working model illustrating how a methylation-phosphorylation crosstalk regulates Nur77 transcriptional activity and the crucial roles of these two modifications in the regulation of EnSC differentiation. Although we performed the experimental treatments to rescue impaired decidualization in RIF by modulating the methylation-phosphorylation of Nur77, further delineation of these molecular mechanisms of the PRMT5/AKT/Nur77 complex will be essential for developing novel therapeutic strategies for embryo implantation failure.

## Methods

### Sex as a biological variable

Our study exclusively examined female mice because the disease modeled is only relevant in females.

### Human endometrial sampling

Endometrial biopsy specimens were obtained from female patients receiving treatment at the Nanjing Drum Tower Hospital’s Centre for Reproductive Medicine. The Control group included women who achieved pregnancy after their first or second IVF-ET treatment for male infertility factor or tubal obstruction. The recurrent implantation failure group (RIF) consisted of patients who had undergone 3 or more consecutive IVF/ICSI-ET or FET cycles, with a cumulative total of at least four high-quality embryos or two high-quality blastocysts that failed to implant. Each woman was required to have regular cyclic menses (25-32 days apart). Exclusion criteria included a known uterine abnormality (e.g., uterine congenital malformation; untreated uterine septum, adenomyosis, or submucous myoma; endometrial polyps; or intrauterine adhesions); a thin endometrium (<6 mm); endometritis diagnosed by hysteroscopy; endometriosis or adenomyosis diagnosed by transvaginal ultrasonography; known autoimmune diseases, currently taking corticosteroids or confounding immunosuppression medications; or abnormal results on parental karyotyping. This study involved a total of 54 endometrial samples from the RIF group and 95 endometrial samples from the control group in the mid-secretory phase. Additional 34 control women were subjected to endometrial biopsies in the proliferative phase, and three others were obtained at 3 days, 5 days and 9 days after hCG administration during natural cycles (referred as LH+3, LH+5 and LH+9). The endometrial biopsies were snap frozen in liquid nitrogen for RNA or protein extraction, or placed in 10% buffered formalin for paraffin embedding, or collected in DMEM-F12 media for isolation of primary endometrial stromal cells. The information of these patients is succinctly presented in Supplemental Table 1.

### Animals and treatments

Uterine stroma-specific mutant mice were generated by crossing *Prmt5^f/f^*mice with Amhr2-Cre mice(25). To conduct the early pregnancy study, C57BL/6 male mice were mated with female *Prmt5^f/f^*and *Amhr2^cre^Prmt5^f/f^* mice at 8 weeks of age. The observation of a vaginal plug marked the beginning of gestation, referred to as day 1 (D 1). To examine implantation, pregnant mice were killed in the morning of D5 to D8. Implantation sites were visualized by an intravenous injection of Chicago blue dye solution, and the number of implantation sites, demarcated by distinct blue bands, was recorded. Mice that failed to recover any embryos were excluded in statistical analysis. To stimulate decidualization in vivo, female mice aged 6-8 weeks with the *Prmt5^f/f^* and *Amhr2^cre^Prmt5^f/f^* genotype were subjected to ovariectomy while receiving suitable pain-relieving medications. Following a period of 14 days, the mice received a subcutaneous injection of 100 ng of estrogen (E2, Sigma-Aldrich, #E2758) for three consecutive days. Following a 2-day period of rest, the mice received a subcutaneous injection of 1 mg of progesterone (P4, Sigma-Aldrich, #P0130) and 10 ng of E2 for three consecutive days. After the most recent hormone injection, one of the uterine horns was subjected to artificial decidualization by injecting 20 μL of sesame oil into its lumen. The other uterine horn was left untreated as a control. Uterine, ovarian, and serum samples were obtained on various days throughout pregnancy.

### Cell culture and in vitro decidualization of endometrial stromal cells

According to the previously mentioned protocol, the primary human EnSCs were extracted and cultivated^13^. Immortal human EnSCs were obtained from ATCC (CRL-4003). To induce decidualization, EnSCs were cultured in DMEM/F12 (Corning, USA) containing 2.5% charcoal/dextran treated FBS (HyClone, USA), 100IU/ml penicillin, and 100 μg/ml streptomycin. The concentrations of 0.5 millimolar (mM) 8-Br-cAMP (Sigma, #B7880) and 1μM medroxyprogesterone acetate (MedChemExpress, #HY-B0469) were applied. The hormonal counselling was denoted as MPA+cAMP. Prior to induce decidualization, cells were exposed to pretreatment with Ad-shPRMT5, Ad-siNur77, Ad-Flag-Nur77, Ad-PRMT5 (adenoviruses produced and stored by the Centre for Molecular Reproductive Medicine, Nanjing University) or peptides for a duration of 48 hours. Cycloheximide (MilliporeSigma, #508739) was introduced at a concentration of 50μg/mL for the specified durations to conduct the protein degradation experiment. For agonist or inhibitor studies, cells were treated with 5nM Cytosporone B (MedChemExpress, #HY-N2148) or 2.5nM MK-2206 (MedChemExpress, #HY-108232) for 48 hours. Subsequently, the cells were exposed to MPA+cAMP for a period of 3 days.

### Construction of adenoviruses

As previously stated, we generated the adenovirus that carries PRMT5 (Ad-PRMT5), shPRMT5 (Ad-shPRMT5)(24), Nur77 (Ad-Flag-Nur77) and siNur77 (Ad-siNur77)(47). The adenoviruses were isolated by CsCl banding and subsequently dialyzed against 20 mmol/L Tris-buffered saline with 10% glycerol after being propagated in HEK293A cells.

### Recombinant protein purification

There were four fusion proteins: GST-Nur77-NT, GST-Nur77-LBD, GST-Nur77-DBD, and GST-Nur77-DBD^R346K^. The fusion proteins were produced in BL21 Escherichia coli bacteria using the pGEX4T-1 vector. To stimulate the expression, a solitary colony was introduced into 5 mL of Luria broth medium supplemented with 100mg/mL ampicillin. Next, 300μl of Pierce Glutathione Agarose (Thermo Scientific, #16101) that had been pre-treated with cold PBS was introduced to the lysate supernatant. The mixture was then incubated at 4℃ for 3 hours with constant mixing. The proteins were subsequently separated using 300μl of elution solution containing 15 mM glutathione, 50 mM Tris-HCl, pH 8.0. Following dialysis in Slide-A-Lyzer Dialysis Cassettes (Thermo Scientific, #66380) using a solution containing 50mM Tris-HCl (pH 7.5) and 10% glycerol at a temperature of 4℃ for a duration of 48 hours, the proteins were separated into groups and subsequently kept at a temperature of −80℃.

### Peptides

The sequences of the examined peptides from Nur77 are as follows: peptide1: NH2-G RLPSKPKQYGRKKRRQRRR-COOH, peptide2: NH2-RTDSLKGRRGRLPSKPKQ YGRKKRRQRRR-COOH, peptide2 mutant: NH2-RTDSLKGRKGRLPSKPKQYGR KKRRQRRR-COOH. All peptides were synthesized (GenScript Biotech) at more than 98% purity, as verified by HPLC and mass spectrometry.

### In vitro methylation

To perform the experiment, 0.5µg of GST-Nur77-NT/DBD/LBD, either wild or mutant type, and recombinant PRMT5/MEP50 (Sigma-Aldrich, #SRP0146) was mixed with 0.5 µg in a buffer solution. The buffer solution consisted of 50mM Tris-HCL (pH 8.6), 2mM MgCl2, 10mM DTT, 0.02% Triton X-100, and 50uM S-adenosylmethionine (Sigma, # A4377). Subsequently, the concoction was left to ferment at ambient temperature for a duration of 2 hours. To stop the reactions, 5×Laemmli sample buffer was added, and the solution was subjected to heating at a temperature of 95°C for a duration of 5 minutes. Ultimately, the proteins underwent analysis via western blotting and Coomassie blue staining (Beyotime, #P0017A) was used for verification.

## Electrophoretic mobility shift analysis (EMSA)

The nuclear extracts of human EnSCs were obtained by use of a nuclear extraction kit (Beyotime, #P0027). GenScript Biotech (Nanjing, China) synthesized and labelled oligonucleotides with Cy5 that matched the binding sequence of Nur77 (NBRE; 5’-GGTAAAGGTCAGGTTGC-3’). The process of binding reactions was carried out utilizing a Lightshift EMSA Optimisation and Control kit (Thermo Scientific, #20148X), in accordance with the guidelines provided by the manufacturer. The binding specificity was assessed using competition studies and the LI-COR Pearl Imaging System was used to perform imaging.

### Immunohistochemistry analysis

The tissues were fixed in 4% paraformaldehyde for 24 hours before being embedded in paraffin wax. The tissue sections underwent deparaffinization and rehydration before being exposed to antigen retrieval. The sections were exposed to the primary antibody (Supplemental Table 2) for an extended time at a temperature of 4℃. This was followed by the immunohistochemical staining kits provided by Zhongshan Golden Bridge. The Leica DM 2000 microscope and LAS Core software (Leica Microsystems Limited, Wetzlar, Germany) were used to take digital images. The protein expression levels in the epithelial cells and stromal cells of the endometrial samples were quantitatively analyzed using the integrated optical density (IOD) of the digital pictures (×400) with the Image-Pro Plus System 6.0 as described previously(45).

### Immunofluorescent staining

The cells underwent fixation by treatment with a 4% paraformaldehyde solution for a duration of 20 minutes. Next, the samples were treated with 0.1% Triton X-100 in PBS to make them permeable for 5 minutes at room temperature. To avoid non-specific binding, the cells were treated with a solution of 3% BSA in PBS and incubated at a temperature of 37℃ for a duration of 1 hour. Subsequently, primary antibodies (Supplemental Table 2) were introduced, and the cells were subjected to overnight incubation at a temperature of 4°C. The signal was visualized using fluorescence-conjugated secondary antibodies. Subsequently, the nuclei were subjected to DAPI staining for a duration of 5 minutes, and images were captured utilizing fluorescence confocal microscopy.

### Co-immunoprecipitation

Tissues or cells were subjected to protein extraction using a lysis buffer consisting of 1% NP-40, 150 mmol/L NaCl, 50 mmol/L Tris (pH 8), 100 μmol/L EDTA, and protease inhibitors. The cell lysates were subjected to pre-clearance using protein A/G beads (Abmart, #A10001M) at a temperature of 4℃ for a duration of 2 hours. Subsequently, 5μg of the primary antibody or isotype IgG was introduced into the purified cell extracts and left to incubate overnight at a temperature of 4℃. Subsequently, protein A/G beads were introduced into the cell extracts and subjected to incubation at a temperature of 4℃ for a duration of 4 hours. For immunoprecipitation of exogenous, Flag-beads (MilliporeSigma, #F1804) or Myc-beads (MilliporeSigma, # E6654) were introduced to the purified cell extracts and left to incubate overnight at a temperature of 4℃. Ultimately, the proteins that were attached to the beads were released by introducing 2xLaemmli sample buffer and subjecting them to a temperature of 95℃ for a duration of 5 minutes. The proteins that were separated and collected were examined using western blotting.

### Western blot analysis

Proteinase and phosphatase inhibitor cocktails (Roche Life Science, #11697498001) and 50 mM Tris-HCl [pH 7.6], 150 mM NaCl, and 1.0% NP-40) were contained in a whole-cell lysis buffer (MilliporeSigma, #P5726) to homogenise tissues and cells. The proteins underwent separation using a 10% SDS-PAGE gel and were subsequently transferred onto PVDF membranes (Millipore, #03010040001). Subsequently, the membranes were subjected to incubation with primary antibodies (Supplemental Table 2), which were subsequently followed by a secondary antibody conjugated with HRP. The detection was carried out via an improved chemiluminescence kit from MilliporeSigma, with the product code #32106. To determine the comparative prevalence of the target proteins, the level of expression for each protein was adjusted based on the level of expression of GAPDH in the corresponding sample. The signal intensities were measured using densitometric analysis with ImageJ software as described previously(45).

### Silver nitrate staining and liquid chromatography–tandem mass spectrometry (LC-MS/MS)

The proteins obtained from co-immunoprecipitation were examined using 10% SDS-PAGE and detected using the Fast Silver Stain Kit (Beyotime, #P0017s) following the directions provided by the manufacturer. Meanwhile, LC-MS/MS was performed by Hoogen Biotech (Shanghai, China). The raw data utilized for proteome profiling were available in Supplemental Table 3.

### RNA isolation and quantitative real-time PCR

Cells underwent TRIzol treatment (Takara Bio, #T9108) for the extraction of total RNA, following the guidelines provided by the manufacturer. The assessment of RNA purity involved quantifying the optical density at wavelengths of 260 nm and 280 nm, whereas the determination of RNA integrity was accomplished through the utilization of agarose gel electrophoresis. 1μg sample of total RNA was used to synthesize the first strand of DNA (cDNA) using the Takara PrimeScript RT reagent kit (Takara Bio, #RR037A). The analysis of gene expression levels was conducted using SYBR Premix Ex Taq kits (Takara Bio, #RR820A) and the corresponding primers (Supplemental Table 4) by real-time PCR. The 2-ΔΔCt method was utilized to compute the relative levels of gene expression, with 18S RNA as the internal control.

### RNA-seq and data analysis

Endometrial stromal cells were cultured in 60 mm dishes and exposed to Ad-shPRMT5 for a duration of 48 hours. Subsequently, they underwent decidualization for an additional 72 hours. The cells were subjected to RNA extraction using TRIzol (Takara Bio, #T9108). The process of RNA sequencing (RNA-seq) was conducted, and the subsequent analysis of the data was carried out by BGI Genomics Co., Ltd, located in Shenzhen, China. The clean reads were aligned to the human reference genome using HISAT2. DESeq2 was utilized to detect differentially expressed genes (DEGs), employing a significance threshold of P value < 0.05 and |log2foldChange| > 1 to determine significant differences. ClusterProfiler was utilized to conduct GO, KEGG, and GSEA analyses. We analyzed the transcription factor regulatory network using KnockTF (http://www.licpathway.net/KnockTF/index.php.)

### Enzyme-linked immunosorbent assay

Following a period of 72 hours of decidualization, the culture supernatants of human EnSC were gathered and subsequently subjected to centrifugation to eliminate cellular debris. The PRL levels were quantified using a commercially accessible enzyme-linked immunosorbent assay kit (R&D Systems, Minneapolis, #DPRL00) in accordance with the instructions provided by the manufacturer. The samples were analyzed twice, and the level of PRL was quantified as nanograms per millilitre of the liquid surrounding the cells.

### Alkaline phosphatase (ALP) staining Assay

The frozen sections were warmed up and fixed for 1 minute following the instructions of the manufacturer (Solarbio, #G1480). ALP incubation was dripped to cover the tissues for 1 hour, followed by nuclear red solution staining. Photographs were captured using a Leica DM 2000 microscope and LAS Core software.

### Luciferase reporter assay

In the pGL3-basic luciferase reporter plasmid (Promega, #E1751), the NBRE synthetic triple sequence repeats were introduced. Sequencing of the plasmid was performed to verify the successful cloning. Undifferentiated human EnSCs at 60% confluency were transfected with the specified plasmids in 24-well plates. After a period of 48 hours, the cells were collected and the luciferase activity was quantified using the Dual-Luciferase Assay System (Promega, #E2940) and a luminescence counter (Berthold Technologies) following the instructions provided by the manufacturer. The activity of Firefly luciferase was adjusted to account for variations in transfection efficiency by normalizing it to the activity of Renilla luciferase.

### Cell viability assays (CCK8 assay)

The experiment involved plating equal numbers of human EnSCs cells in 96-well plates. The cells were then treated with Ad-shPRMT5, and incubated for 12h, 24h, 36h, or 48h. Cell viability was assessed via a Cell Counting Kit-8 (CCK8) assay, in accordance with the instructions provided by the manufacturer. The optical density at 450nm (OD450nm) was then measured.

### Three-dimensional structure prediction of protein-protein interaction

To investigate the binding regions and interaction patterns among PRMT5, Nur77, and AKT proteins, we employed the professional protein-protein and protein-DNA/RNA docking program HDCOK. The structure with the highest docking score was selected as the standard result for subsequent interaction analysis. The docking scores were based on the ITScorePP or ITScorePR iterative scoring functions (59, 60).

### Statistical analysis

The experiments were conducted a minimum of three times. The statistical analyses were conducted using Prism version 9 software developed by GraphPad, or R software. The data shows the biological replicates’ mean ± SEM. A two-tailed Student’s t-test was employed to compare the average expression values between the two treatment groups. A one-way ANOVA was conducted to compare many groups. Two-way analysis of variance (ANOVA) with the Bonferroni multiple comparisons test was performed to analyze the interaction effects of more than two groups.

### Study approval

The Institutional Review Boards at Nanjing Drum Tower Hospital granted approval for the human research (2013-408081-01). All patients provided informed consent prior to the sampling procedure. The Institutional Animal Care and Use Committee of Nanjing Drum Tower Hospital (20210510) granted approval for all animal experiments.

## Data availability

The authors provide detailed description of methods and original data upon request. RNA-seq data sets generated in this study have been deposited at the NCBI database with BioProject accession number PRJNA1050378.

## Author contributions

GY, RJ, HS, and XC initiated and supervised the project. XC, RJ, ZC and XW, performed the experiments, and collected the data. YL, XT and XZ contributed to the animal models and animal analysis. MW, NK and LD contributed to the human endometrium and endometrial stromal cell experiments; RJ, and XC wrote the manuscript. GY, HS, and JS reviewed and edited the manuscript.

## Acknowledgements

This work was supported by the National Natural Science Foundation of China (82030040 to H.S., 82171653 and 82371680 to G.Y., 82271698 to R.J. and 82301899 to X.C.), the National Key Research and Development Program of China (2023YFC2705402 to G.Y).

## Notes

### Competing Interest Statement

The authors have declared no competing interest.

